# Gene Delivery Mediated by Backbone-Degradable RAFT Copolymers

**DOI:** 10.1101/2025.10.17.682953

**Authors:** Prajakatta B. Mulay, D. Christopher Radford, Brayan Rondon, Bruna Favetta, Benjamin S. Schuster, Jia Niu, Adam J. Gormley

## Abstract

Cationic polymers present an attractive platform for gene delivery. However, these highly charged macromolecules can also lead to cytotoxicity. Therefore, there is a strong unmet need to develop efficacious polymeric gene delivery vehicles with high biocompatibility. Here, we leveraged recent advances in polymer chemistry to develop backbone-degradable cationic copolymers and evaluate their potential as gene delivery vehicles. Specifically, polycations were prepared via copolymerization with macrocyclic allylic sulfides which can participate in PET-RAFT polymerization via radical ring-opening cascade copolymerization to install degradable backbone segments. A polymer library with varying degradability was prepared and evaluated using a model GFP plasmid to transfect U-2 OS cells. Incorporation of degradable groups into the copolymer backbone improved transfection efficiency 10-fold at low amine/phosphate (N/P) ratios without increasing cytotoxicity, thereby enhancing their value as gene delivery carriers. We hypothesize that degradability may enhance the complex’s disassembly kinetics in the cytosol, enabling more efficient payload release.

## INTRODUCTION

Gene therapy presents a promising alternative to traditional therapeutics in treating hereditary as well as non-hereditary diseases including genetic disorders, neurological disorders, cardiovascular diseases, and cancer.^1^ To do this, DNA and RNA are used to correct, modify, or silence missing genes to treat diseases.^2,3^ However, the application of gene therapy faces challenges due to the fragile nature of these therapeutic genes, which are susceptible to degradation by serum nucleases, have poor membrane permeability, low cellular uptake, and exhibit poor stability in circulation.^4–6^ Therefore, gene delivery vectors are critical for delivering these therapeutic genes into the target cells and ensuring efficient transfection. These vectors are divided into two main types: viral and non-viral. Viral vectors, despite their efficiency, pose safety risks such as immune responses and toxicity.^7^ Non-viral vectors, which include cationic polymers, lipids, and nanoparticles, are preferred for their lower immunogenicity, cost-effectiveness, high loading capacity, and versatility.^8–10^ Among these non-viral vectors, cationic polymers are highly versatile, exhibit batch-to-batch uniformity, and possess reasonable control over their macromolecular structure.^11^ Common examples of cationic polymers for gene delivery include polyethylenimines (PEI), poly(2-N-(dimethylaminoethyl) methacrylate) (PDMAEMA), and poly(L-lysine) (PLL).^12–14^ Cationic polymers can condense with negatively charged nucleic acids via electrostatic interactions to form polyelectrolyte complexes also known as polyplexes. These polyplexes are taken up by cells through endocytosis followed by the endosomal release in the cytosol, where they can disassemble to release their genetic payloads and traffic to their intracellular site of action.^15^ For example, polyplexes formed with plasmid DNA (pDNA) must translocate to the nucleus for transcription and protein expression.^16^

The main challenge for the clinical application of cationic polymers is cytotoxicity, arising primarily due to their high molecular weight, positive charges, and often non-degradable nature.^17^ This concern over biocompatibility is further exacerbated by the need for repeated administrations with many gene therapies that can cause downstream issues with accumulation and clearance.^9,18^ While using lower molecular weight polymers can mitigate toxicity, it also decreases their ability to complex with therapeutic genes and transfect cells.^19–22^ This trade-off complicates the design and use of polyplexes for gene delivery, creating the need for a dynamic gene delivery vector. Biodegradable polymers have the potential to address these challenges by degrading inside the lysosome thus lowering their accumulation in treated cells and overall toxicity.^9,23–25^ The degradation of biodegradable polymers in physiological environments relies on hydrolysis of the polymer backbone via breakdown of labile linkages such as esters. This process allows these degradation products to be safely eliminated from the body through excretion, improving their biocompatibility. Biodegradability may also enhance disassembly kinetics of the polyplexes in the cytosol enabling more efficient payload release.^26, 27^ The first examples of backbone degradable polymers for gene therapy were explored by Park and coworkers with poly(4-hydroxy-L-proline ester) (PHP).^28^ They found that PHP degrades to half its original molecular weight in under two hours and fully degrades in three months, showing effective pDNA binding and comparable transfection efficiency to PLL. Similarly, cationic polylactides also demonstrated successful gene transfection with complete hydrolytic degradation within one week.^29^ Although multiple investigations have been conducted to evaluate the transfection efficiency of backbone degradable polyesters, there is a lack of evidence of their degradation kinetics.^30–36^

Alternatively, vinyl polymers have been widely used in gene delivery applications due to their synthetic versatility and straightforward synthesis.^11^ Specifically, controlled/living free radical polymerization strategies such as reversible addition-fragmentation chain transfer (RAFT) enable the synthesis of well-defined vinyl polymers with narrow dispersity from a diverse library of monomers that can be tailored for different applications.^37^ Such advantages have previously proven useful in generating cationic polymer libraries for subsequent evaluation as synthetic gene delivery vehicles.^11^ For example, previous work by Reineke and coworkers has identified copolymers of AEMAm and HEMA as an efficacious delivery platform for such efforts.^38,39^

However, vinyl polymers are inherently non-degradable, as the chemistry is limited to backbones consisting exclusively of carbon-carbon bonds.^40^ This, in turn, can lead to biocompatibility concerns for this class of polymer. To overcome this challenge, various radical ring-opening polymerization (rROP) techniques have been explored.^41^ In these strategies, cyclic monomers are introduced to the reaction to copolymerize with the vinyl monomers, enabling incorporation of hetero-atoms into the otherwise all-carbon backbone. This provides a straightforward means to install labile chemical groups (e.g., esters, thioesters, disulfides, etc.) that facilitate biodegradability. However, early rROP monomer candidates such as cyclic ketene acetals (CKAs) and thionolactones demonstrated numerous unfavorable properties including poor incorporation, unbalanced reactivity ratios with the vinyl comonomers, ring-retaining side-reactions that fail to introduce degradability, and failure to copolymerize with certain classes of vinyl monomers.^41^ While earlier studies have demonstrated some success with leveraging these systems for improved gene delivery^42–44^, the application of backbone-degradable RAFT copolymers in this field remains underexplored.

To address the limitations of these cyclic monomers, Niu and coworkers recently developed macrocyclic allylic sulfone monomers that are able to participate in the RAFT process via radical ring-opening cascade copolymerization (rROCCP).^45^ These monomers demonstrated broad compatibility with a variety of vinyl co-monomers and near-unity reactivity ratios with acrylate and acrylamide co-monomers.^45,46^ Importantly, these monomers were also shown to be compatible with oxygen-tolerant photoinduced electron/energy transfer RAFT (PET-RAFT) polymerization chemistry. PET-RAFT has recently enabled controlled polymerizations to be performed on the bench-top at room temperature using milder conditions in addition to offering temporal control.^47,48^ PET-RAFT polymerizations can be performed in a well-plate format which enables generating a library of diverse copolymers that can be coupled with automation for accelerated discovery of gene-delivery vehicles.^49–51^ In addition, such mild conditions also allow the synthesis of cationic RAFT copolymers^52–54^, that can lead to facile incorporation of degradable monomer units which may otherwise degrade during thermal-initiated polymerizations.

Herein, we build upon this work by leveraging rROCCP to install labile ester groups directly into the backbone of cationic copolymers. We hypothesized that incorporation of backbone degradability into this system would improve their function as gene delivery vectors, enhancing biocompatibility and transfection efficiency. Towards this end, we first confirmed ester groups incorporated by this chemistry imparted biodegradability to the resulting polymer products. Importantly, chain fragmentation was observed in response to enzymatic challenge in an esterase-rich environment simulating the lysosome. We then applied this chemistry to generate backbone-degradable cationic copolymers via PET-RAFT polymerization, copolymerizing a cationic monomer, a hydrophilic monomer, and a macrocyclic allylic sulfide to form backbone-degradable copolymers (Fig. 1A). A variable number of biodegradable residues were incorporated into the backbone to create four distinct backbone-degradable copolymers. The ester groups introduced in these residues enable hydrolytic degradation of the copolymers and fragmentation of the polymer chain (Fig. 1B). The backbone-degradable copolymers were then complexed with a model GFP-encoded plasmid to evaluate their transfection efficiency in vitro in U-2 OS cell line which is a commercially available cell line conducive for transfections (Fig. 1C). Finally, the cytotoxicity induced by these degradable copolymers was also investigated to identify the top-performing backbone-degradable polyplexes. These results demonstrate a promising proof-of-concept that biodegradable polyplexes can be synthesized using PET-RAFT chemistry, paving the way for improved designs.

**Figure 1.**
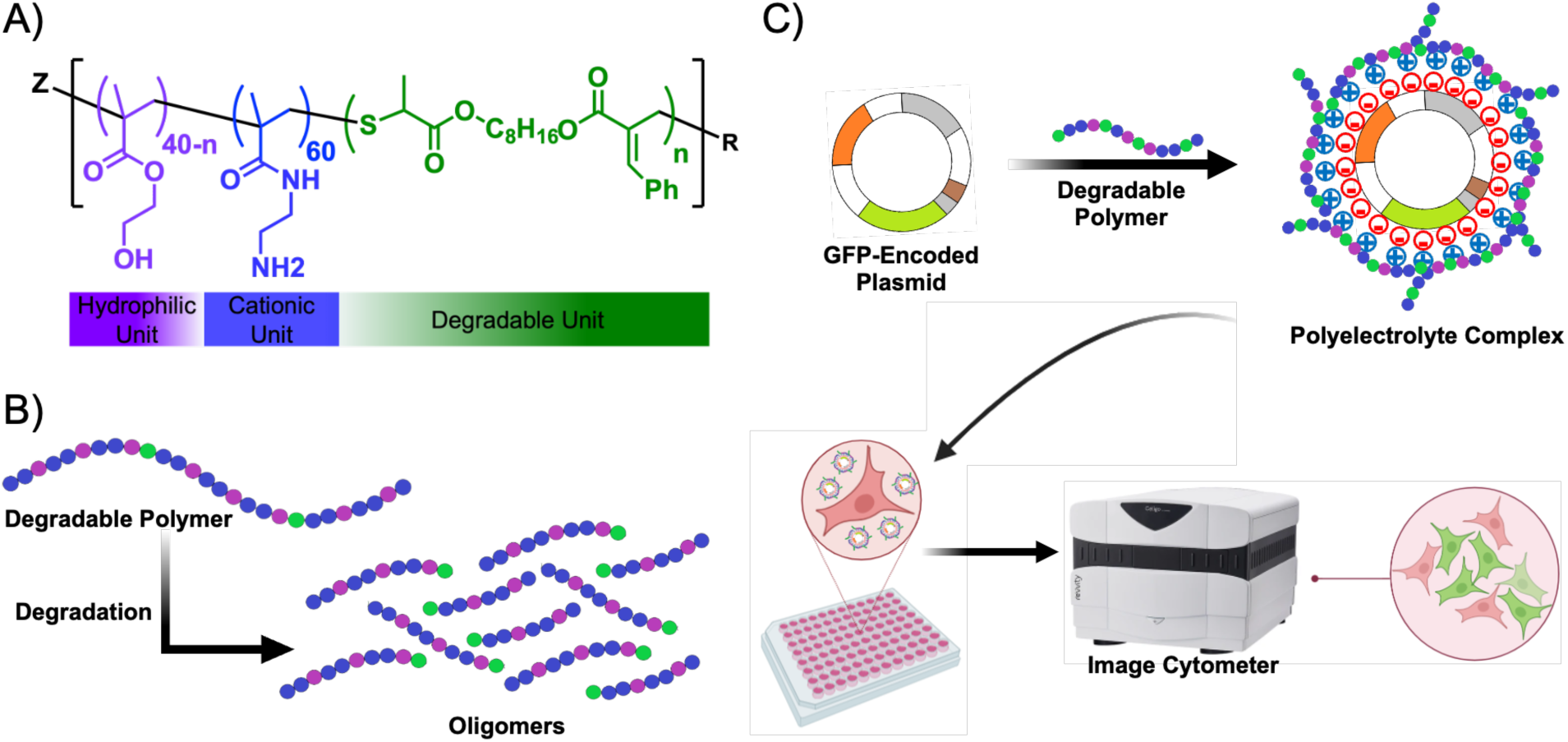
Synthesis and evaluation of degradable polyplexes. (A) Chemical structure of the backbone-degradable copolymer containing hydrophilic (2-hydroxyethyl methacrylate, HEMA), cationic (2-aminoethyl methacrylamide, AEMAm), and macrocyclic allylic sulfide monomer (Cyc1). (B) Ester groups in the degradable copolymer backbone undergo hydrolytic degradation. (C) Degradable copolymers form a polyplex upon condensation with a GFP-encoded plasmid (pMAX-GFP) via electrostatic interactions. U-2 OS cells were transfected with the polyplexes formed with backbone-degradable polymers. GFP-positive cells were counted using an image cytometer.

## MATERIALS AND METHODS

### Materials

All reagents were obtained from Sigma-Aldrich unless otherwise stated. Macrocyclic allylic sulfide monomer Cyc1 was synthesized as previously described.^45^ The U-2 OS cell line (RRID:CVCL_0042) was obtained from American Type Culture Collection (ATCC, Catalog No. HTB-96).

### Polymer synthesis and characterization

#### General procedure

Polymers were synthesized via PET-RAFT using established reaction conditions.^47^ In brief, stabilized monomers were first de-inhibited before use by passing over MEHQ inhibitor removal resin. Stock solutions of chain transfer agent (CTA, 50 mM), photoinitiator tris(2-phenylpyridinato-C^2^,*N*)iridium(III) (*fac*-Ir(ppy)_3_, 1 mM), and requisite monomers were prepared in DMSO. These stocks were then used to prepare the reaction solutions. Photopolymerization was then carried out under 450 nm LED light at room temperature to yield the polymer product. Monomer conversions were determined using ^1^H-NMR spectroscopy (Bruker Avance Neo 500 MHz) of crude product with mesitylene as an internal standard. Crude products were optionally purified by precipitation into hexane:acetone (10:1) mixture three times and dried under vacuum. Monomer content of purified polymers was analyzed by ^1^H-NMR in D_2_O. A typical procedure for PET-RAFT copolymerization of each polymer are as follows:

#### poly(DMA)

Dimethylacrylamide (DMA) was co-polymerized with Cyc1 using 2-(2-carboxyethylsulfanylthiocarbonylsulfanyl)propionic acid as CTA. The reaction was carried out a final concentration of 2 M total monomer, 4 mM CTA, and 50 µM *fac*-Ir(ppy)_3_, with 2.5 mol% Cyc1 in the monomer feed. The reaction mixture was irradiated for 6h to yield the polymer product. Non-degradable poly(DMA) control was prepared in parallel by analogous reaction conditions, with the exception of excluding Cyc1 from the reaction.

#### Cationic copolymers

2-aminoethyl methacrylamide (AEMAm) and 2-hydroxyethyl methacrylate (HEMA) were co-polymerized with Cyc1 using 4-cyano-4-(((ethylthio)carbonothioyl)thio)pentanoic acid as CTA. Reactions were carried out at a final concentration of 1 M total monomer, 10 mM CTA, and 200 µM *fac*-Ir(ppy)_3_. AEMAm feed ratio was fixed at 60 mol% for all reactions, with Cyc1 feed ratio varying from 0-10 mol%, and HEMA filling the balance. The reaction mixture was irradiated for 18 h to yield the polymer product.

### Polymer degradation

#### Chemical degradation

To assess degradation of polymers in response to chemical challenge, polymer reaction mixtures were diluted 100-fold into 50 mM NH_4_OH and the solution was incubated for 30 mins at 37°C. The solution was then diluted 2:1 in dimethylformamide (DMF) and molecular weight of the polymer products were then analyzed via size exclusion chromatography (SEC) using an Agilent 1200 Series system with on-line UV and RI (Agilent 1260 Series) detectors. The system was equipped with two Agilent PLgel 5 μm columns in series (10^3^ and 10^4^ Å, 300 x 7.5 mm). DMF supplemented with 50 mM LiBr was used as the mobile phase. Molecular weight data (*M_n_*, *M_w_*, and *Đ*) were determined using a series of PMMA standards of known molecular weight (Agilent EasyVial PEG Calibration Kit) based on the respective RI chromatographs.

#### Enzymatic degradation

Enzymatic degradation was carried out using porcine liver esterase (PLE) as a model enzyme. Fresh solutions of PLE (Sigma Aldrich) were prepared from lyophilized powder at 20 U/mL in 10 mM HEPES buffer (pH 7.0). Prior to use, activity of esterase solution was confirmed via chromogenic assay, with 4-nitrophenyl butyrate serving as substrate. Polymer reaction mixtures were then diluted 100-fold into the enzyme solution and incubated for 24 or 48h at 37°C using a shaker-heater set to 400 RPM. At the appropriate time point, the solution was removed from the heater-shaker and diluted 2:1 in DMF to precipitate the enzyme. The solution was then centrifuged (15k RCF, 5 min, RT) to pellet protein precipitate, and the supernatant was analyzed via SEC on the same instrumentation described above. A corresponding 0 h time point was prepared by adding polymer to the enzyme solution and immediately precipitating the enzyme. Hydrolytic degradation of the polymer in the absence was esterase was assessed using an analogous procedure.

### Polyplex formation

A shuttle vector, pMAX_GFP was obtained from Addgene (plasmid# 177825). Gene sequences were verified by whole plasmid sequencing (Azenta). Plasmids were transformed into DH5α competent *E. coli* (New England Biolabs). Colonies picked from fresh plates were grown for 12 hours at 37°C in 5 mL LB while shaking at 250 rpm. The vectors used contained a kanamycin resistance gene; kanamycin was used at concentrations of 50 μg/mL in cultures. pDNA was extracted using a Monarch® Plasmid DNA miniprep kit (New England Biolabs) using the manufacturer’s protocol. The extracted pDNA was stored at-20°C in aliquots in an elution buffer. The DNA concentration in each aliquot was measured based on its absorbance at 280 nm using a Nanodrop spectrophotometer (ThermoFisher). One aliquot of pMAX_GFP was then diluted in sterile filtered DNase-free water to a concentration of 20 ng/µL. The copolymers were also diluted in sterile filtered DNase-free water to the desired concentration and mixed in equal volumes with pMAX_GFP to obtain the desired N/P ratios of 5, 10, and 20. N/P ratio is the stoichiometric ratio between the protonable nitrogen (N) in the copolymer and the anionic phosphate groups (P) present in the pMAX_GFP. These ratios were chosen based on the previously reported recommendations for transfection with cationic RAFT copolymers.^38^ The polyplex was then incubated for 45 minutes at room temperature and then mixed with two parts volume of serum-free DMEM and incubated for an additional 45 minutes at room temperature. Polyplexes were also prepared with PEIpro (Polyplus) as the commercial transfecting agent following the manufacturer’s protocol. Briefly, equal volumes of pMAX_GFP (40 ng/µL) and PEIpro solution (80 µL/mL) were mixed in Opti-MEM reduced serum medium (ThermoFisher Scientific) and incubated at room temperature for 45 minutes.

### Cellular Assays

The U-2 OS cell line was used to assess the transfection efficiency where cells were maintained in DMEM supplemented with 10% FBS at 37°C and 5% CO_2_ in 75cm^2^ cell culture flasks. For all transfection assays, cells were seeded at 6000 cells/well at 200 µL/well in a tissue culture-treated 96-well plate and incubated at 37°C and 5% CO2 for 24 h before transfection. The transfection protocol was adapted from Kumar, et. al.^38^ Media was aspirated after 24 h and a 200 ng/well plasmid loading was employed where cells were incubated with 60 μL of the polyplex-serum-free-DMEM solution for 4 h at 37°C and 5% CO2. After 4 h, 200 μL DMEM with 10% FBS was added to all the wells. For transfection with PEIpro, 24 h after seeding, media was aspirated, and cells were supplemented with growth medium. After 4 h, 10 µL of PEIpro-pMAX_GFP polyplex was added to the growth medium-supplemented cells. Media was aspirated from all the wells after 24 h, supplementing the cells with 200 µL of fresh growth medium. The GFP expression, cell counts, and cell viability were evaluated at 48h after transfection. All treatments were performed in triplicate.

Cell viability was performed first using a CCK-8 assay (Dojindo) according to the manufacturer’s protocol. At 44 h after transfection, 20 µL of 2% solution of CCK-8 was added to the cells and incubated for a total of 4 h at 37°C and 5% CO2. Absorbance values were obtained every hour at 450 nm using a SpectraMax UV-Vis plate reader. Blank values obtained in empty wells containing only the media and CCK-8 solution were subtracted from all measurements. Absorbance values were normalized to the control cells (cells treated with 60 µL of DNase-free water and serum-free DMEM solution for 4h) to determine the cell viability.

GFP expression was then immediately measured using target expression analysis with a Celigo Image Cytometer (Nexcelom Bioscience). Green fluorescence channel (483/536 nm) was used to image the GFP expression with an exposure time of 10 ms. Celigo software was used for the automated image analysis that counts the GFP-positive cells and the mean intensity of GFP expression in each cell. After GFP expression analysis, the cells were then stained with Hoechst 33342 (ThermoFisher Scientific), a widely used dye for live cell imaging that stains the nucleus blue. The GFP expression and the Hoechst stain were not imaged simultaneously to reduce the interference caused by the overlap of the green and blue channels. Hoechst staining was carried out according to the manufacturer’s protocol to determine the total number of cells in each well. Briefly, media was aspirated from all the wells and stained with 30 µL of the Hoechst solution (prepared by diluting 1:2000 in D-PBS) for 10 minutes. After 10 minutes, the cells were washed three times with 100 µL PBS and were imaged in PBS as well. Target expression analysis was again performed using the Celigo Image Cytometer with a blue (377/447 nm) channel with an exposure time of 300 ms. Celigo software was used for the automated image analysis that counts the Hoechst-stained cells.

### Statistical Analysis

All data are reported as mean ± standard error. The number of replicates per experimental and control group is three as also described in the related Methods subsections and the figure captions. Statistical analysis of all data was processed using the Origin software package (OriginPro, USA). Statistical significance between degradable and non-degradable RAFT copolymers datasets at respective N/P ratios was determined using Student’s t-test (two-tailed with equal variances). Data was normalized to the highest observed value for plotting transfection efficiency and GFP cell count. For cell viability, data was normalized to the control group of cells.

## RESULTS AND DISCUSSION

### Biodegradable Polymers

The macrocyclic allylic sulfide Cyc1 was utilized as a comonomer to directly introduce ester groups into the polymer backbone during the polymerization process (Figure S1). Building on previous work by Niu^45^, poly(DMA) was selected as an initial water-soluble polymer system to investigate hydrolytic degradability of these esters. Towards this end, a copolymer of DMA and Cyc1 (2.5 mol% feed ratio) was synthesized via PET-RAFT polymerization. A DMA homopolymer was also synthesized using identical reaction conditions to create a control polymer lacking the backbone ester groups. Both polymers presented similar molecular weights (*M*_n_ values of 44.6 kDa vs 42.6 kDa, respectively) and dispersities (*Ɖ* values of 1.38 vs 1.40, respectively), indicating that introduction of esters via rROCCP did not compromise reaction control of the PET-RAFT chemistry, consistent with previous reports.^45^

The polymer pair was then subjected to chemical and enzymatic challenge to assess hybridizability of ester groups, with changes in molecular weight analyzed by size exclusion chromatography (SEC, Figure 2a). Upon incubation with 50 mM NH_4_OH for 30 mins, a significant shift in the SEC profile was observed (*M*_n_ = 10.2 kDa), as well as a corresponding increase in dispersity (*Ɖ* = 1.73) (Figure 2b, red traces). This indicated the ester bonds in the backbone were rapidly hydrolyzed, leading to fragmentation of the polymer chain. In contrast, no change in molecular weight was observed for the DMA homopolymer lacking the ester residues in its backbone (Figure 2b, black traces).

**Figure 2.**
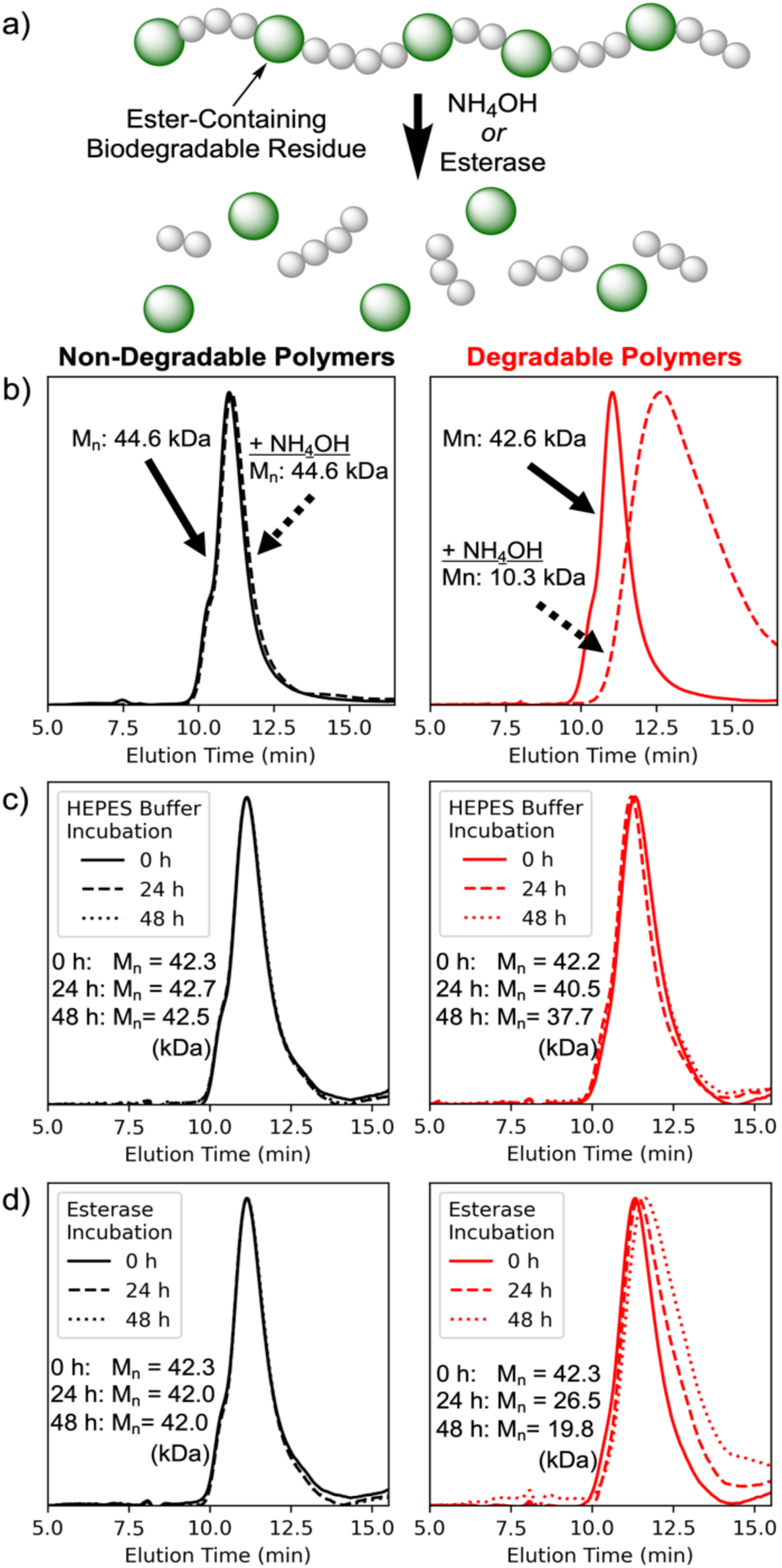
Biodegradability of copolymers. a) Copolymerization with Cyc1 monomer incorporates ester-containing biodegradable residues into the polymer backbone. Size-exclusion chromatography (SEC) was used to analyze polymers after incubation with b) NH_4_OH, c) HEPES buffer, and d) esterase. Polymers synthesized without the macrocyclic co-monomer (black traces) show no change in molecular weight. In contrast, ester-containing copolymers (red traces) saw significant shifts in response to chemical and enzymatic challenge, while remaining stable under neutral conditions.

Based on these promising results, biodegradability of the polymers was then assessed using a model esterase (porcine liver esterase). Importantly, the ester-containing copolymer demonstrated hydrolytic stability in the absence of enzyme, with minimal shift in the SEC trace over 48h (Figure 2c). However, when incubated with esterase, a gradual degradation of the polymer chain was observed (Figure 2d, red traces). As expected, esterase had no effect on the molecular weight of the homopolymer (black traces), demonstrating that the ester groups incorporated by rROCCP could serve as sites of enzymatic degradation. Such stimuli-sensitive degradability has clear utility for gene delivery vectors, as it can allow robust extracellular stability, while facilitating rapid disassembly and release of the nucleic acid payload upon endocytosis and trafficking to the esterase-rich lysosomal environment.

### Biodegradable Cationic Polymers

Building upon this promising result, we next investigated whether this chemistry could be applied to polymer designs suitable for use as synthetic gene delivery vehicles. Specifically, a design based on an AEMAm-HEMA copolymer was selected as a model system due to previous reports of its success in this role.^38, 39^

A PET-RAFT polymerization was conducted with Cyc1 (10 mol% feed ratio), AEMAm (60 mol% feed ratio), and HEMA (30 mol% feed ratio) monomers to synthesize a biodegradable cationic polymer with a target degree of polymerization (DP) of 100 (Scheme 1). Time points were collected throughout the reaction and ^1^H NMR was used to independently monitor reaction kinetics of the three monomers over the course of the polymerization (Figure 3a). The Cyc1 monomer was observed to convert less efficiently than the other monomers (only reaching 61.5% conversion after 18h versus >99% and 89.7% for HEMA and AEMAm, respectively. However, when comparing Cyc1 conversion versus total monomer conversion (Figure 3b), the macrocyclic monomer continues to undergo conversion throughout the course of the reaction, indicating ester groups are being incorporated throughout the length of the growing polymer chain. This distribution should facilitate significant fragmentation of the backbone upon degradation and could, in turn, improve payload release.

**Figure 3.**
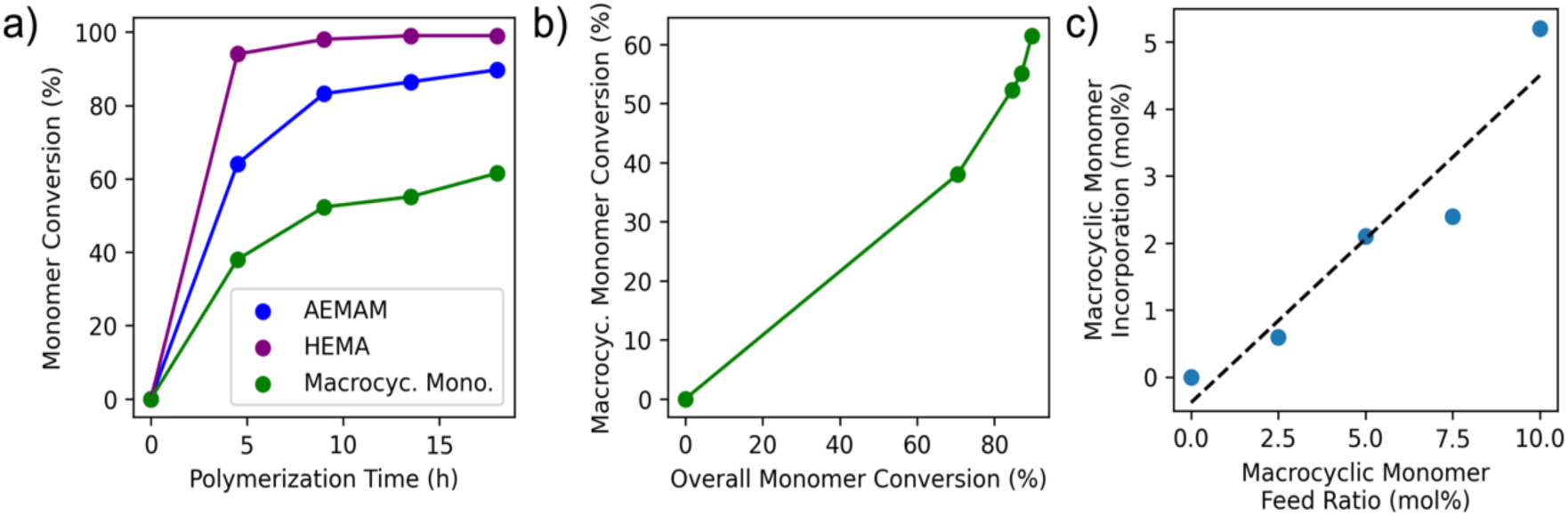
Polymerization kinetics and incorporation of macrocyclic allylic sulfide monomer (Cyc1) into backbone-degradable cationic copolymers via rROCCP. a) Copolymerization kinetics of a PET-RAFT reaction polymerizing co-monomers AEMAm (60 mol% feed ratio), HEMA (30 mol%), and Cyc1 (10 mol%). Monomer conversion is presented as percent of its respective feed. b) Cyc1 conversion as a function of overall monomer conversion across the kinetics study. Conversion continues to increase over the course of the reaction, indicating incorporation of the degradable ester sequence throughout the polymer chain. c) Comparison of degradable sequence frequency in copolymer backbone across five PET-RAFT polymerization reactions with varying Cyc1 feed ratio (0-10 mol%), showing degradability can be modulated by altering monomer feed.

**Scheme 1.**
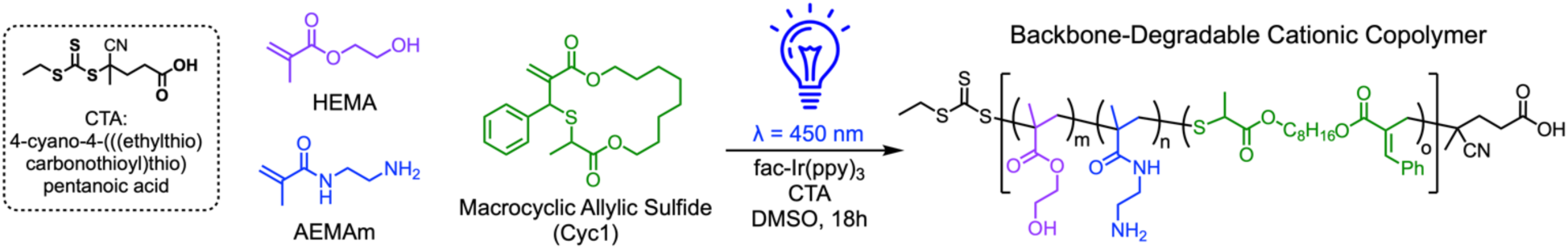
Synthetic scheme for synthesis of backbone-degradable cationic copolymers via PETRAFT polymerization. Macrocyclic allylic sulfide Cyc1 can participate in PET-RAFT via radical ring-opening cascade copolymerization (rROCCP) to introduce ester groups into the copolymer backbone.

Next, a series of five PET-RAFT polymerizations of cationic copolymers were conducted, all targeting a theoretical DP of 100, but varying Cyc1 feed ratios in order to assess the relationship between monomer feed ratio and content of the degradable sequence in the copolymer product. Cationic comonomer content was held constant in the monomer feed (60 mol%), with Cyc1 and HEMA being varied from 0–10 mol% and 30–40 mol% respectively. ^1^H NMR analysis of the copolymer products (Figures S2-S6) indicated a clear linear relationship between feed ratio of the Cyc1 monomer and frequency of the degradable sequence in the backbone (Figure 3c), suggesting that the degradability can be conveniently tuned simply by altering the feed ratio of the Cyc1 monomer.

### Polyplex Formation and Transfection with Backbone-Degradable Cationic Copolymers

A library of cationic AEMAm-HEMA copolymers (Figures S7-11 and Table S1) containing variable numbers of biodegradable residues were then evaluated for their ability to complex the model GFP plasmid (pMAX_GFP) and successfully transfect U-2 OS cells. Transfection efficiency was quantified using high-throughput cell imaging using a green fluorescence channel (483/536 nm) at a range of N/P ratios of 5, 10, and 20. N/P ratio is the stoichiometric ratio between the protonable nitrogen (N) in the copolymer and the anionic phosphate groups (P) present in the pMAX_GFP. Copolymers of AEMAm and HEMA at a DP of 100 have previously been shown to successfully delivery GFP gene payloads and induce significant expression.^38, 55^ Therefore, we varied the feed ratio of Cyc1 within this copolymer scaffold design (60% AEMAm, 0–10% Cyc1, 30–40 % HEMA) to establish that moderate backbone-degradability could be introduced without significant loss of function. Figure 4a shows the transfection efficiency obtained with the polyplexes formed with degradable copolymers. The transfection efficiency was calculated as the number of GFP-expressing cells divided by the total number of cells (obtained by Hoechst staining). It can be observed from Figure 4a that complexes formed with high Cyc1 feed ratios of 7.5% and 10% demonstrate the highest transfection efficiency at low N/P ratios (i.e., low polymer content) of 5 and 10. However, at a higher N/P ratio of 20, i.e. at the highest copolymer concentration, their transfection efficiency significantly drops, owing to increased cytotoxicity (discussed further in the following section). At lower Cyc1 content of 2.5 and 5 mol%, substantial transfection can still be observed at a high N/P ratio of 20. Complexes formed without Cyc1 (i.e., non-backbone-degradable polymers) demonstrated a linear trend in transfection efficiency with increasing N/P ratio; however, their transfection was significantly lower than complexes formed with Cyc1 at low N/P ratios of 5 and 10 as observed in Figure 4a. Specifically at N/P ratio of 5, a 10-fold increase in transfection is observed with complexes containing 7.5% and 10% Cyc1 compared to their non-backbone degradable analog. In sharp contrast, untreated cells and those treated with only the pMAX-GFP vector (i.e., no polymer) did not demonstrate any transfection (Figure S12). This suggests that introducing backbone degradability could enhance the disassembly of the polyplex in the cytosol enabling more efficient payload release. Figure 4b demonstrates the actual count of GFP-positive cells observed in each well. Complexes formed from copolymers with high Cyc1 content (feed ratios of 7.5% and 10%) at a low N/P ratio of 5 again express more than twice the number of GFP-positive cells than that observed for polyplexes with lower Cyc1 content. However, at a slightly higher N/P ratio of 10, a lower Cyc1 content of 5% demonstrates higher GFP-positive cells than Cyc1 content of 7.5% and 10% which may be attributed negligible cell death as discussed in the next section. Further increasing the N/P ratio to 20 resulted in reduced GFP expression across all polyplexes as increasing Cyc1 content was not able to rescue expression at high polymer concentrations that can cause higher cellular toxicity. To investigate the amount of GFP molecules expressed in a single cell, the mean GFP intensity obtained from the Celigo image analysis software was plotted for the polyplex treated cells. No significant differences were observed in the intensity between the transfecting agents as observed in Figure 4c except for Cyc1 content of 5% at N/P of 10 that demonstrated slightly higher intensity than the non-degradable copolymers at the same N/P. Therefore, it can be concluded that the ability of the polyplexes formed with degradable copolymers to produce GFP molecules in a cell is comparable to that of the commercial transfecting agent PEIpro.

**Figure 4.**
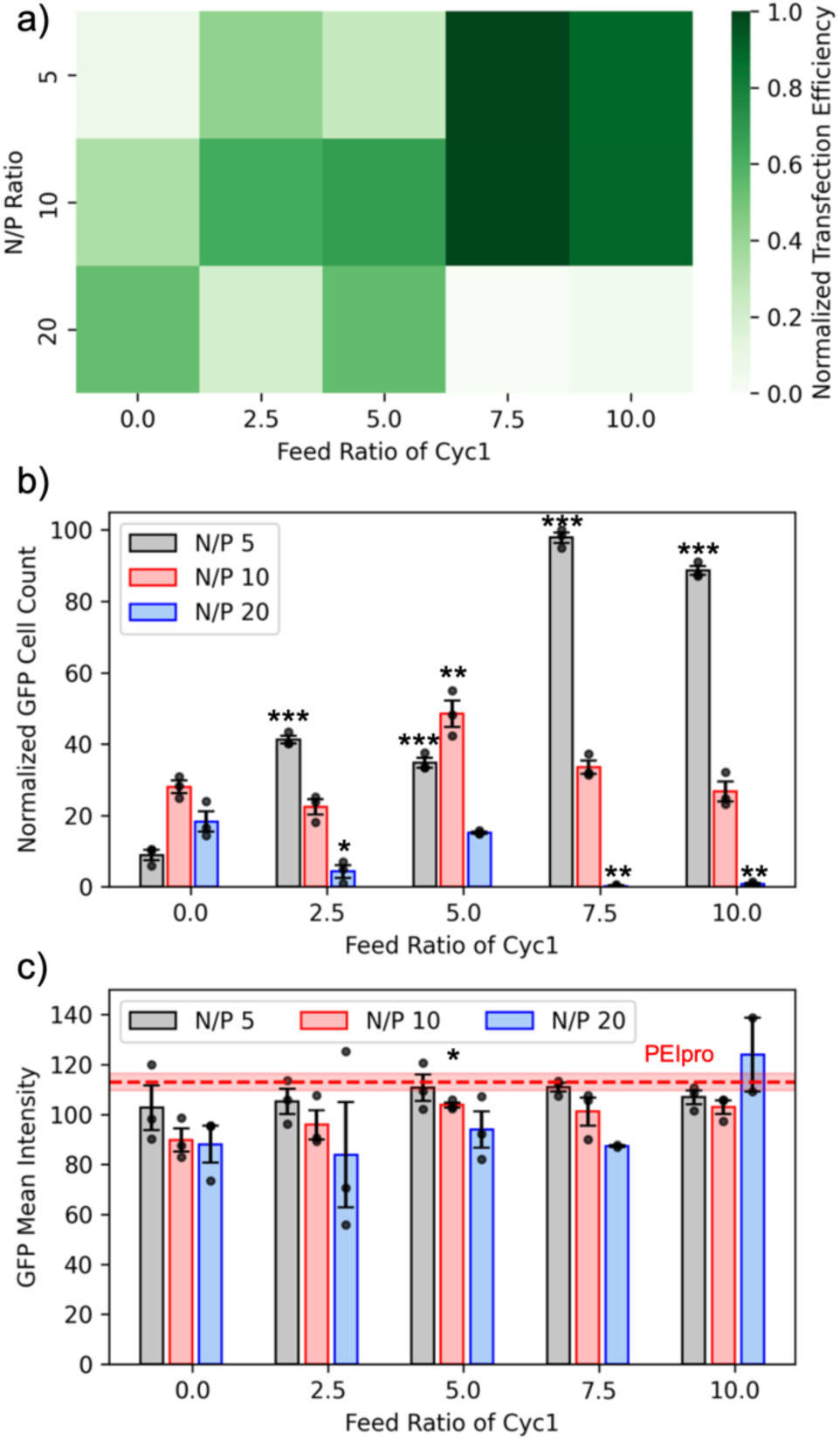
Transfection efficiency of degradable and non-degradable polyplexes. Polyplexes were formed with pMAX_GFP at varied N/P ratios using degradable copolymers with variable macrocyclic biodegradable residues. U-2 OS cells were transfected with these polyplexes and were imaged at 48 h. a) The highest transfection efficiency was observed with degradable copolymers containing 7.5% macrocyclic monomer content at low N/P ratios of 5 and 10. b) The number of cells expressing GFP was high with degradable copolymers containing 7.5% and 10% of macrocyclic monomer at a low N/P ratio of 5. c) The GFP mean intensity, or the average number of GFP molecules produced per cell by polyplex and commercial reagent PEIpro demonstrate no significant difference. Significance between degradable (feed ratio of macrocyclic monomer > 0) and non-degradable RAFT copolymers datasets at respective N/P ratios were determined using Student’s t-test (* p<0.05; ** p<0.01; *** p<0.001). All plots are plotted with mean ± SE at n = 3 replicates.

### Cytotoxicity Evaluation of Polyplex Library

Next, we sought to assess cell death caused by the complexes formed with degradable copolymers. Although evaluating transfection efficiency is an important metric for deciding the best-performing transfecting agent, the fact that it is measured based on the total number of live cells is often overlooked. That is if a transfecting agent is cytotoxic but can transfect as many cells as a non-cytotoxic transfecting agent, the cytotoxic agent will demonstrate higher efficiency due to the reduction in the total number of live cells. Therefore, assessing the cell death caused by transfecting agents is an important metric in deciding the best-performing transfecting agent. We assessed the cell death caused by the degradable copolymers by counting the number of live cells at 48 h after transfection and compared it to the control cells. Figure 5a demonstrates the cell death (%) which indicates that polyplexes formulated at a low N/P ratio of 5 demonstrate less than 20% cell death. The complex formed with 5% Cyc1 content at N/P ratio 5 did not demonstrate any cell death, that is, this formulation led to greater cell proliferation than the control cells. Complexes formed with 7.5% and 10% Cyc1 content at an N/P ratio of 10 had demonstrated high transfection efficiency (Figure 4a); however, it caused nearly 70% cell death, and therefore, its performance comes at the cost of unacceptable cytotoxicity. All the complexes formed at a high N/P ratio of 20, regardless of whether they are degradable or non-degradable, cause nearly 70-80% death compared to the untreated control cells suggesting higher polymer content is toxic to the cells. Even PEIpro with high transfection efficiency causes a nearly 40% reduction in cell count or cell death leading to an artificial increase in the transfection efficiency. Along with degradability, Cyc1 also introduces hydrophobicity to the degradable copolymers due to its structure containing an eight-carbon aliphatic saturated chain. The effects of hydrophobic modification on cationic polymers are controversial and studies have reported negative effects on the cytotoxicity of the gene delivery vector.^56, 57^ In cationic RAFT copolymers, introducing hydrophobicity in the side chain is reported to elevate cytotoxicity.^58^ The increased cytotoxicity for degradable polyplexes with 7.5% and 10% Cyc1 content at N/P ratio of 10 may be attributed to the additional hydrophobicity Cyc1 introduced into the backbone. However, such effect is shown to be mitigated at the lower N/P ratios which also yield the highest transfection efficiency. The viability of the cells was also assessed using a commercial CCK-8 kit. All polyplexes formed at a lower N/P ratio of 5 demonstrated nearly 100% cell viability which is higher than that demonstrated by PEIpro (75%) as shown in Figure 5b. However, at higher N/P ratios of 10 and 20, the cell viability drops below 50% for most complexes formed with degradable copolymers. This demonstrates that polyplexes formed with higher Cyc1 content of 7.5% and 10% and a lower N/P ratio of 5 demonstrate higher transfection efficiency than their non-degradable analog while maintaining low cell death and high cell viability (Figure 6). This suggests that our degradable copolymers have the potential to significantly improve biocompatibility while enhancing gene delivery function.

**Figure 5.**
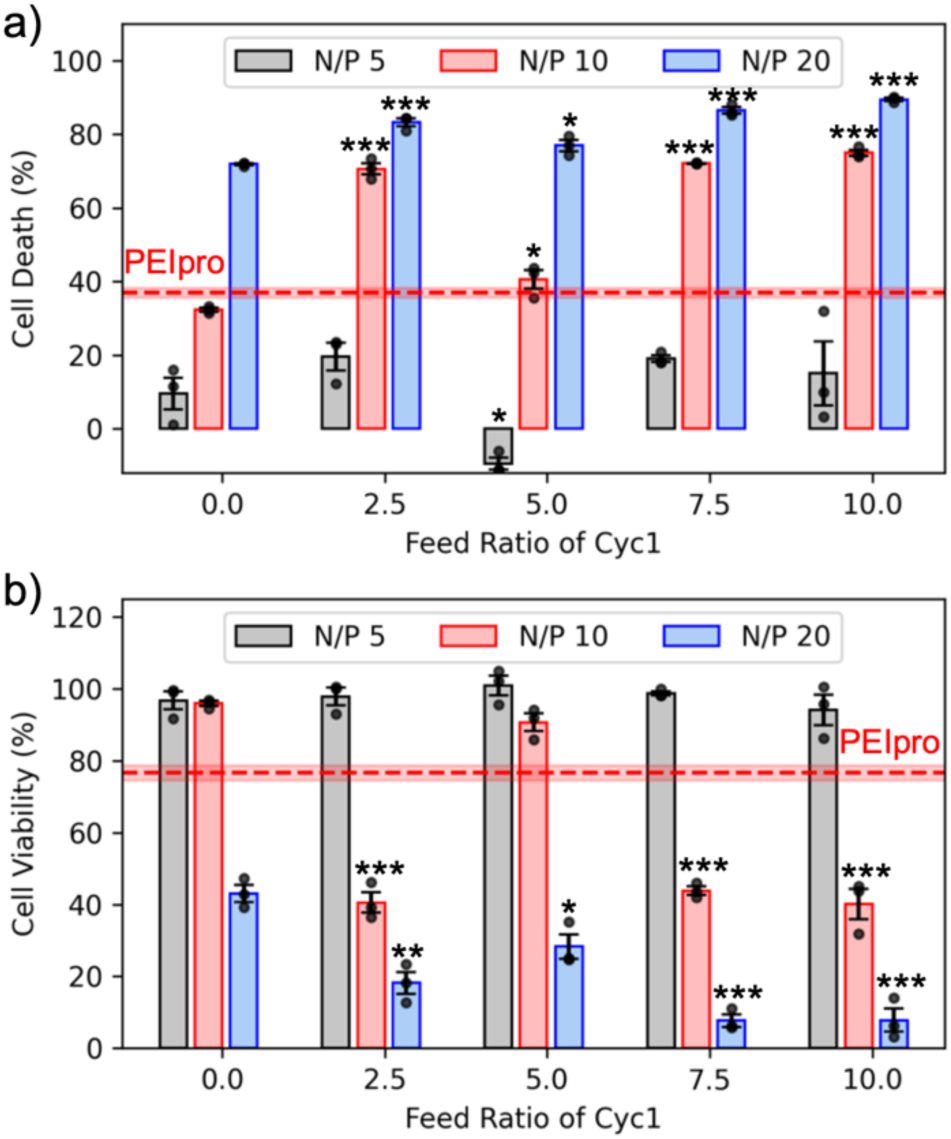
Cytotoxicity of degradable and non-degradable polyplexes. a) Cell death caused by polyplexes formed with pMAX_GFP at varied N/P ratios using degradable copolymers with variable macrocyclic biodegradable residues. Cell death (%) was calculated as [(Y-X)*100/Y] where Y is the total number of cells in the control wells and X is the total number of cells in treated wells obtained by Hoechst stain counting. Higher N/P ratios of 20 causes higher cell death. Commercial reagent PEIpro causes higher cell death than polyplexes formed at N/P 5. b) Cell viability of polyplex treated cells as measured by CCK-8 kit and normalized to the control cells. All polyplexes formulated with degradable copolymers with an N/P ratio of 5 demonstrate comparable cell viability to the control cells and higher cell viability than PEIpro. Significance between degradable (feed ratio of macrocyclic monomer > 0) and non-degradable RAFT copolymers datasets at respective N/P ratios were determined using Student’s t-test (* p<0.05; ** p<0.01; *** p<0.001). All plots are plotted with mean ± SE at n = 3 replicates.

**Figure 6.**
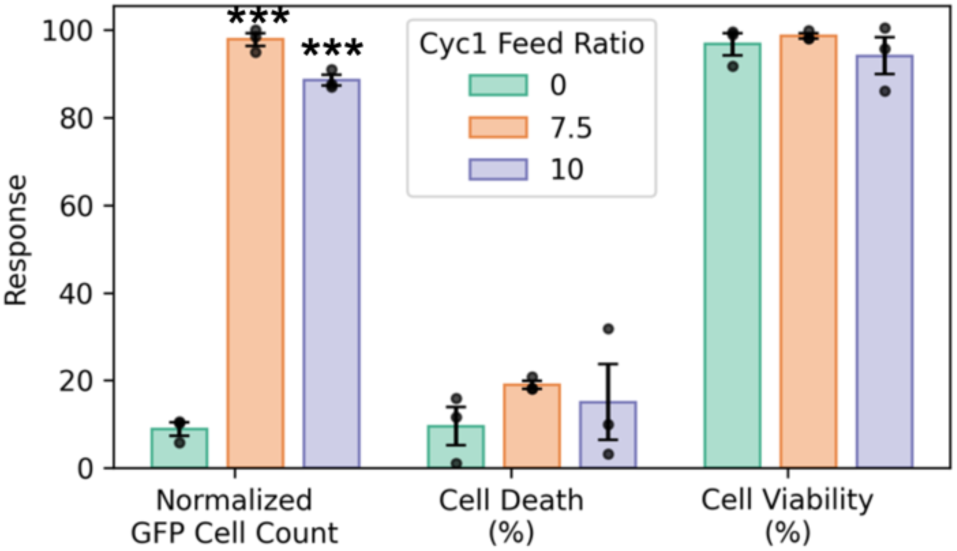
Normalized GFP cell count, cell death, and cell viability obtained for polyplexes formed with degradable copolymers containing 7.5% and 10% Cyc1 content at N/P 5 compared to its nondegradable analog containing 0% Cyc1. Polyplexes formed with 7.5% and 10% Cyc1 content demonstrate higher GFP cell count while maintaining low cell death and high cell viability. Significance between degradable (feed ratio of macrocyclic monomer > 0) and non-degradable RAFT copolymers datasets at respective N/P ratios were determined using Student’s t-test (*p<0.05; ** p<0.01; *** p<0.001). All plots are plotted with mean ± SE at n = 3 replicates.

## CONCLUSION

Synthetic gene delivery vectors based on cationic polymers present an appealing alternative to viral approaches. Polymerization techniques such as PET-RAFT produce well-defined polymer products under mild conditions and can leverage a diverse monomer library to provide a broad chemical space for the development of such therapies. However, traditional vinyl-based copolymers are inherently non-degradable. This, combined with the significant cationic character necessary to complex the genetic payload, raises biocompatibility concerns for these systems.

Herein, we have copolymerized a macrocyclic allylic sulfide to incorporate ester groups into the copolymer backbone of cationic vinyl copolymers. Importantly, because degradability was introduced through a co-monomer rather than bespoke CTAs or post-polymerization modifications, this approach allowed the degree of degradability to be modulated by altering the feed ratio of the macrocyclic monomer. The introduction of these ester groups into the backbone produced copolymers that were able to hydrolytically degrade into lower molecular weight fragments. This approach is particularly attractive for gene delivery applications, as lysosomal conditions could lead to accelerated degradation of ester linkages. This could in turn allow robust stability of the polyplex extracellularly while facilitating rapid disassembly and release of the nucleic acid payload upon endocytosis and lysosomal trafficking.

Using this synthetic strategy, a series of backbone degradable, cationic copolymers were synthesized, varying the feed ratio of the macrocyclic monomer. These copolymers were then used to complex a model GFP plasmid and generate a library of polyplexes with varying N/P ratios. Evaluating this library, biodegradability was shown to significantly improve transfection efficiency, particularly at low N/P ratios (e.g., N/P = 5). Furthermore, copolymers with the highest frequency of degradable residues (7.5 and 10 mol% feed ratios) were the top performers, indicating a clear relationship between the extent of degradability and transfection efficiency. Importantly, this improvement did not come at the expense of biocompatibility. At an N/P ratio of 5, top-performing polyplexes yielded 10-fold higher GFP-positive cells versus non-degradable analogs, while maintaining low cell death and high viability. Collectively, this demonstrated the potential of these degradable copolymers to serve as potent, biocompatible synthetic gene delivery systems.

## ASSOCIATED CONTENT

**Supporting Information**. Additional characterization information for the polymerization chemistry and backbone-degradable polycation library, as well as representative microscopy images demonstrating successful gene delivery using a model GFP plasmid (PDF).

## ACKNOWLEDGMENTS

We would like to thank Dr. Biju Parekkadan for providing access to use the Celigo Image Cytometer.

