## Supporting Information for "Gene Delivery Mediated by Backbone-Degradable RAFT Copolymers"

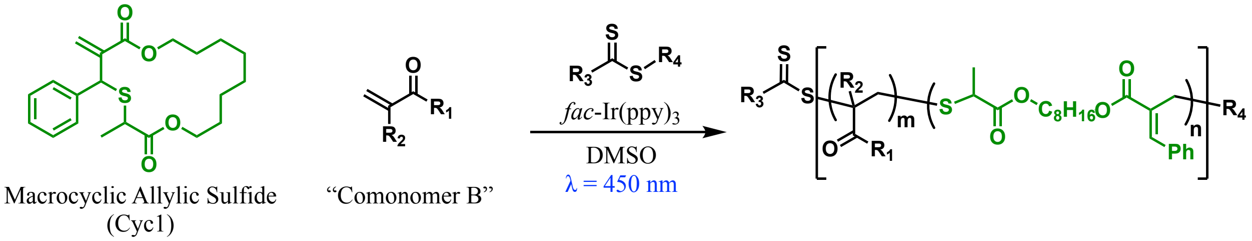


**Figure S1.** General synthetic scheme for copolymerization of macrocyclic allylic sulfide monomer Cyc1 with comonomer (*i.e.*, (meth)acrylates or (meth)acrylamides) via PET-RAFT polymerization. Cyc1 is able to participate in the PET-RAFT process via radical ring-opening cascade copolymerization (rROCCP).

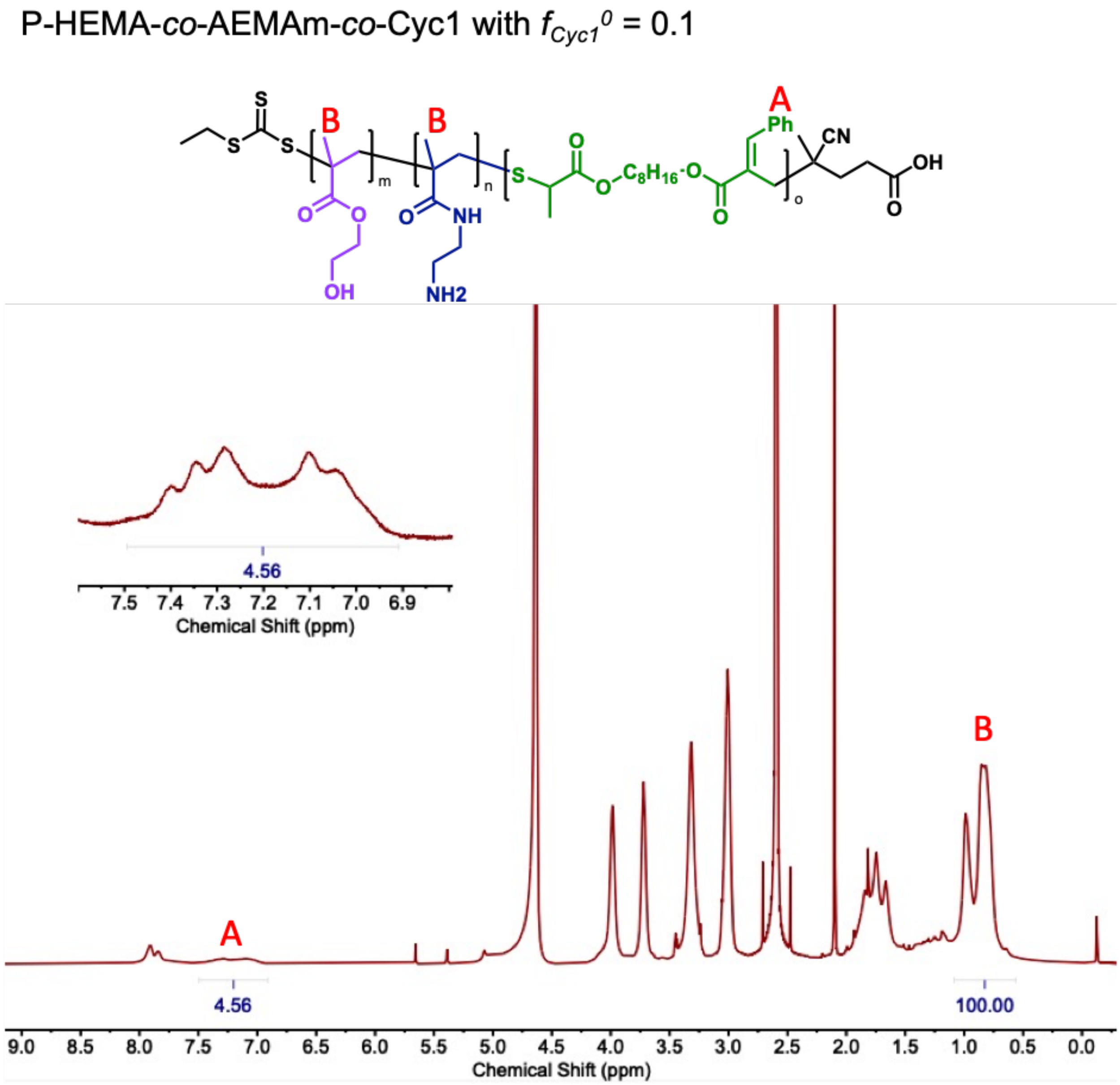


**Figure S2.** $^1$H NMR in $D_2O$ of degradable cationic copolymer prepared with Cyc1 feed ratio of 10 mol%. Degradable unit incorporation calculated by the integration of aromatic protons of the degradable comonomer Cyc1 (A) and methyl protons of HEMA and AEMAm comonomers (B): % incorp. = $[I_a/5]/([I_a/5] + [I_b/6])$ x 100%. As shown in the figure, % incorp. = (0.912/17.578) × 100% = 5.2%.

P-HEMA-*co*-AEMAm-*co*-Cyc1 with $f_{Cyc1}^{0} = 0.075$

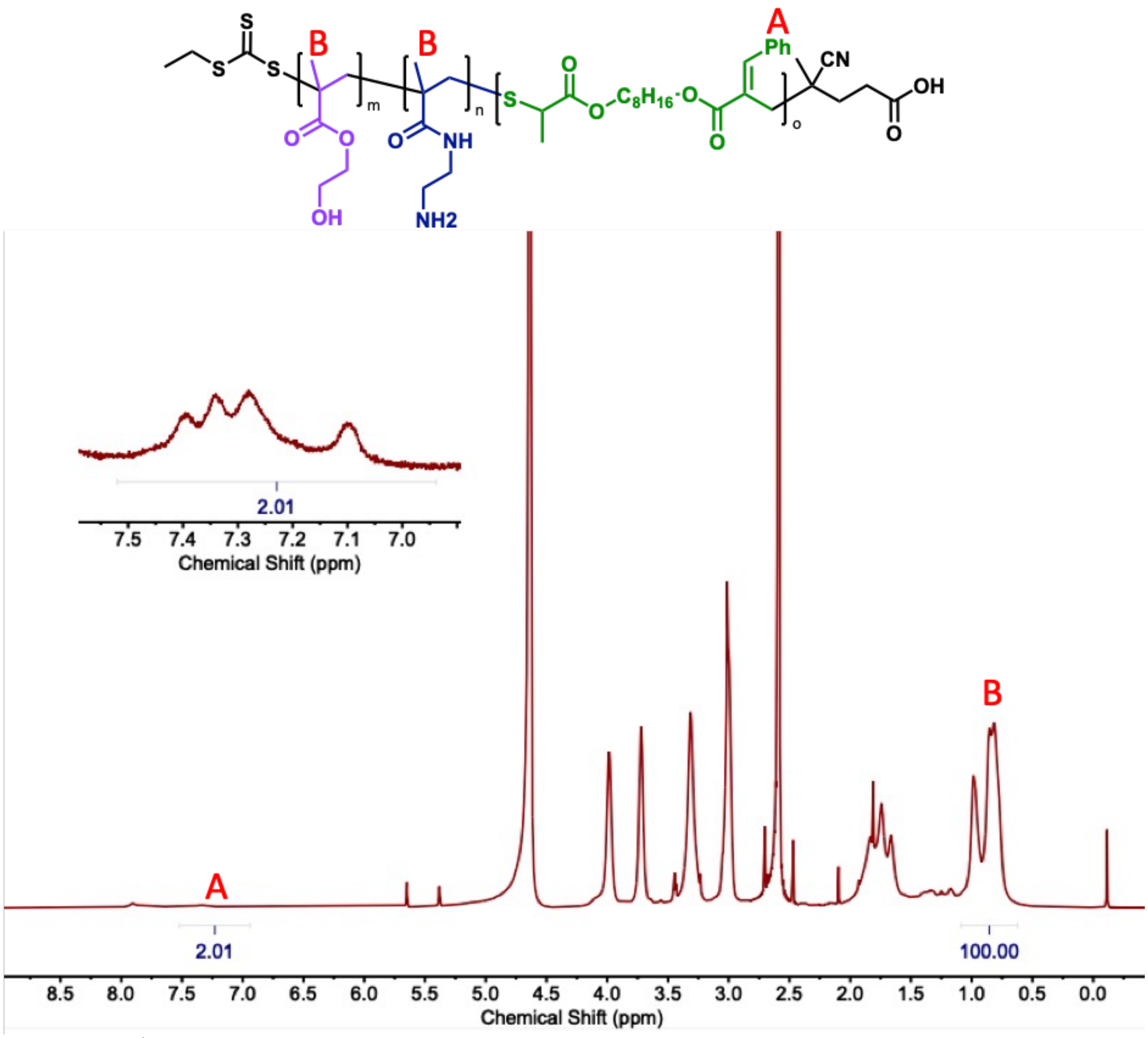


**Figure S3.** $^{1}H$ NMR in $D_2O$ of degradable cationic copolymer prepared with Cyc1 feed ratio of 7.5 mol%. Degradable unit incorporation calculated by the integration of aromatic protons of the degradable comonomer Cyc1 (A) and methyl protons of HEMA and AEMAm comonomers (B): % incorp. = $[I_a/5]/([I_a/5] + [I_b/6])$ x 100%. As shown in the figure, % incorp. = (0.402/17.068) × 100% = 2.4%.

P-HEMA-*co*-AEMAm-*co*-Cyc1 with $f_{Cyc1}^{0} = 0.05$

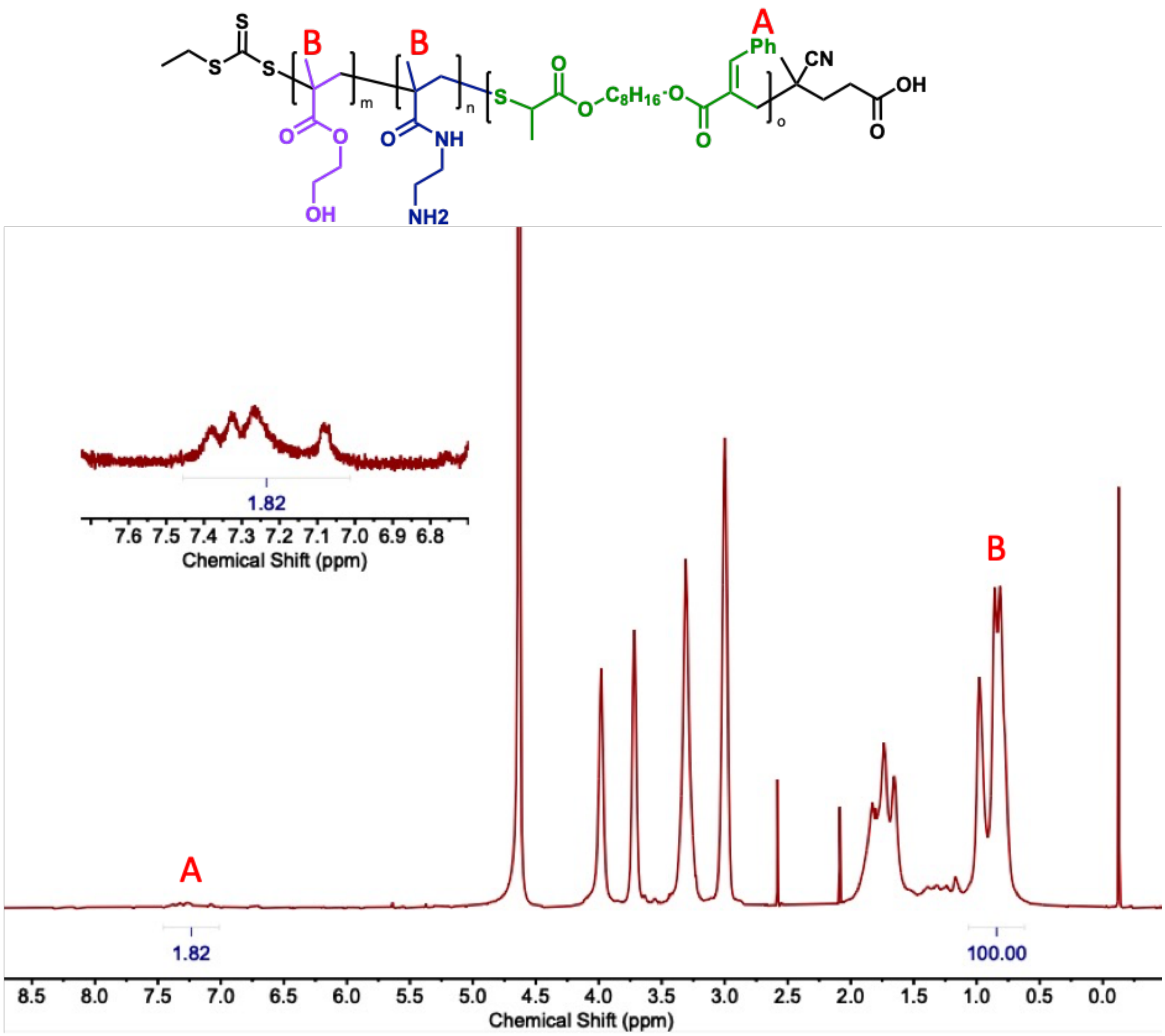


**Figure S4.** $^{1}$H NMR in $D_2O$ of degradable cationic copolymer prepared with Cyc1 feed ratio of 5 mol%. Degradable unit incorporation calculated by the integration of aromatic protons of the degradable comonomer Cyc1 (A) and methyl protons of HEMA and AEMAm comonomers (B): % incorp. = $[I_a/5]/([I_a/5] + [I_b/6])$ x 100%. As shown in the figure, % incorp. = (0.364/17.030) × 100% = 2.1%.

P-HEMA-*co*-AEMAm-*co*-Cyc1 with $f_{Cyc1}^{0}$ = 0.025

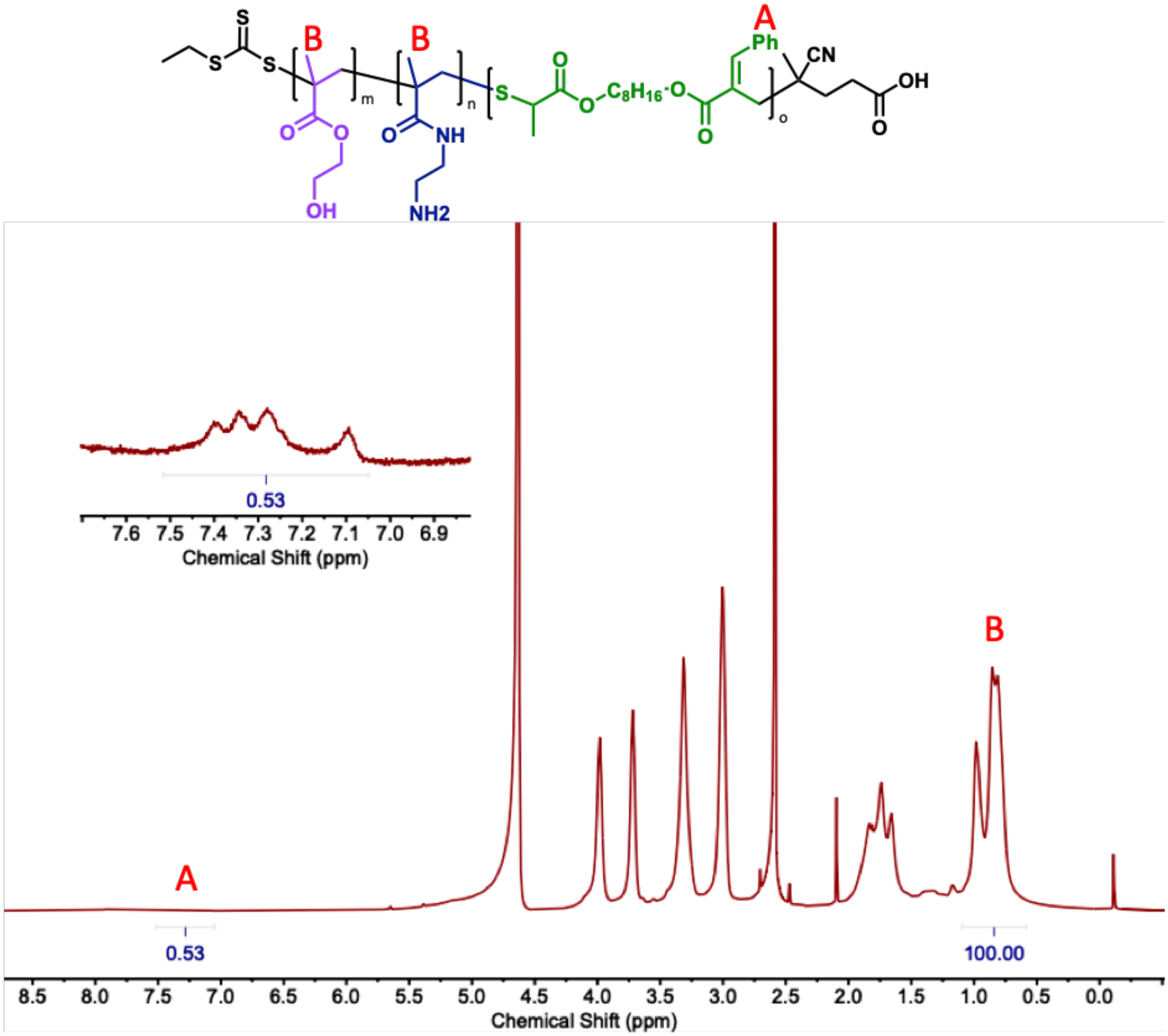


**Figure S5.** $^{1}H$ NMR in $D_2O$ of degradable cationic copolymer prepared with Cyc1 feed ratio of 2.5 mol%. Degradable unit incorporation calculated by the integration of aromatic protons of the degradable comonomer Cyc1 (A) and methyl protons of HEMA and AEMAm comonomers (B): % incorp. = $[I_a/5]/([I_a/5] + [I_b/6])$ x 100%. As shown in the figure, % incorp. = (0.106/16.772) × 100% = 0.6%.

P-HEMA-*co*-AEMAm ($f_{Cyc1}{}^{0}$ = 0)

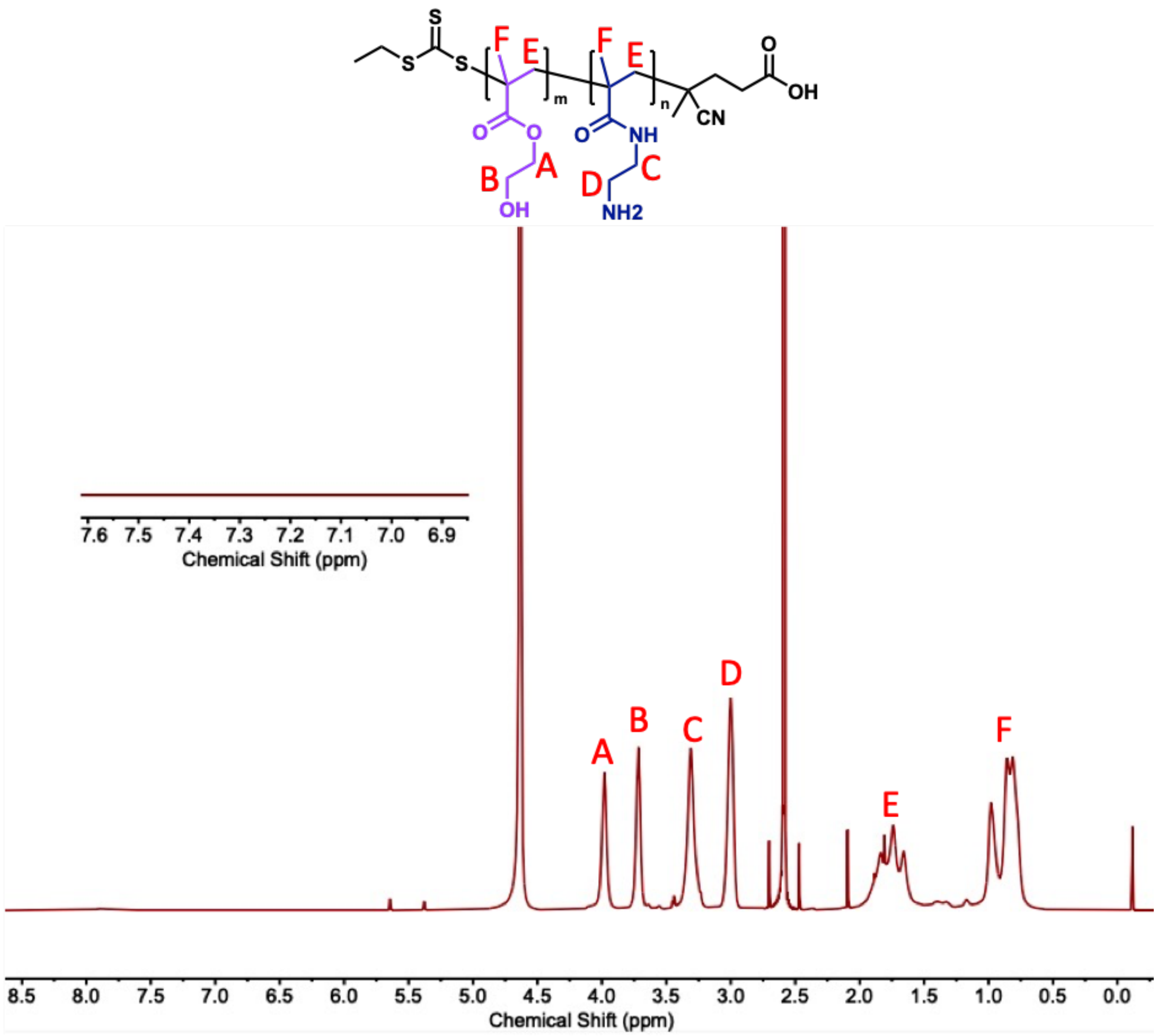


**Figure S6.** $^{1}$H NMR in $D_2O$ of non-degradable cationic copolymer prepared without Cyc1 in feed ratio.

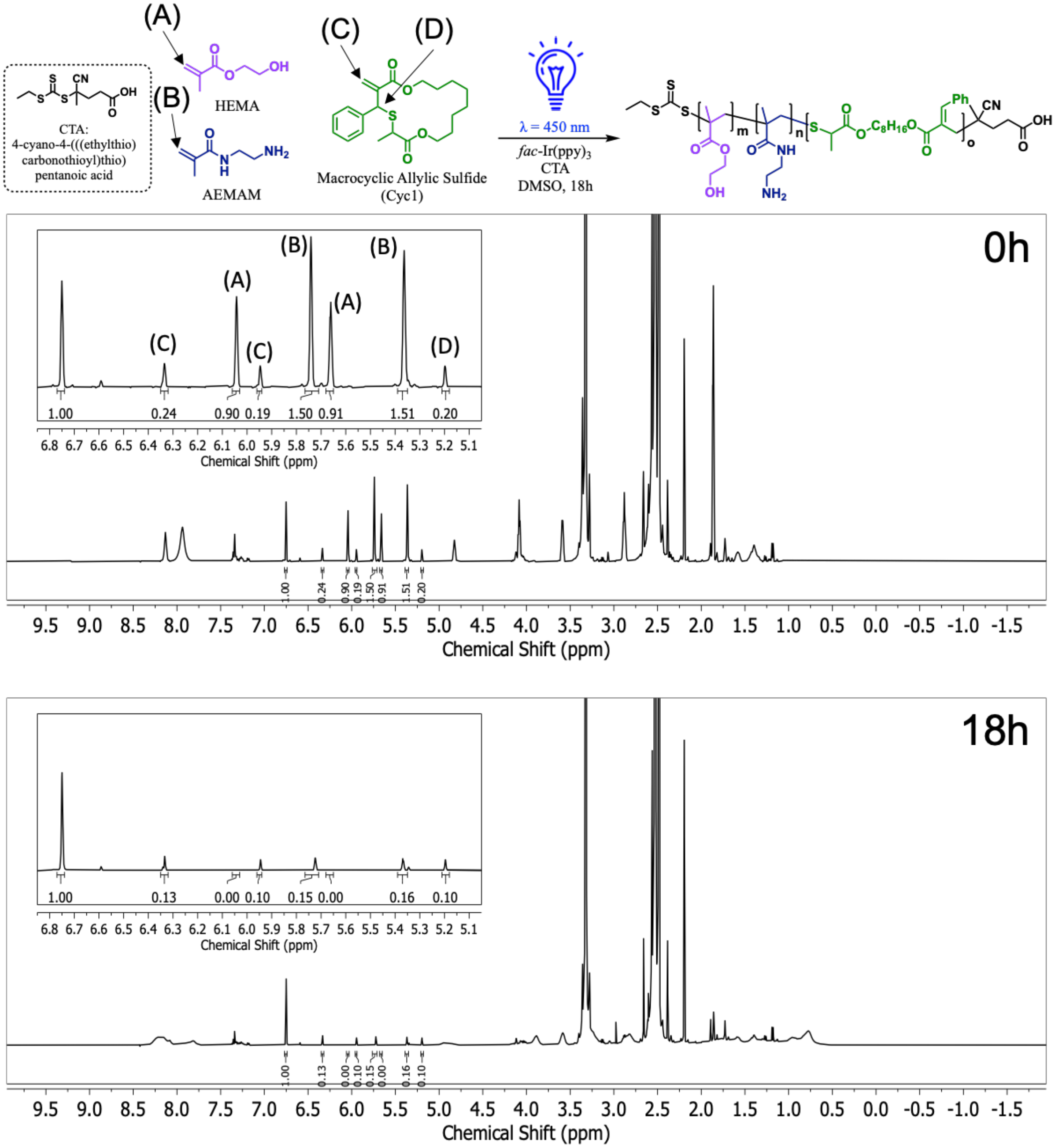


**Figure S7.** [1]H NMR (DMSO-d$_6$) of reaction mixture for degradable cationic copolymer **P1** ($f_{AEMAm}{}^{0}$ = 0.6, $f_{HEMA}{}^{0} = 0.3$, $f_{Cyc1}{}^{0} = 0.1$) at the start (0h, top) and end (18h, bottom) of the reaction. Integrals are assigned to vinyl protons of unreacted HEMA (δ = 6.04 ppm and δ = 5.66 ppm, "A"), vinyl protons of unreacted AEMAm (δ = 5.74 ppm and δ = 5.36 ppm, "B"), vinyl protons of unreacted Cyc1 (δ = 6.34 ppm and δ = 5.95 ppm, "C"), and allylic proton of unreacted Cyc1 (δ = 5.20, "D").

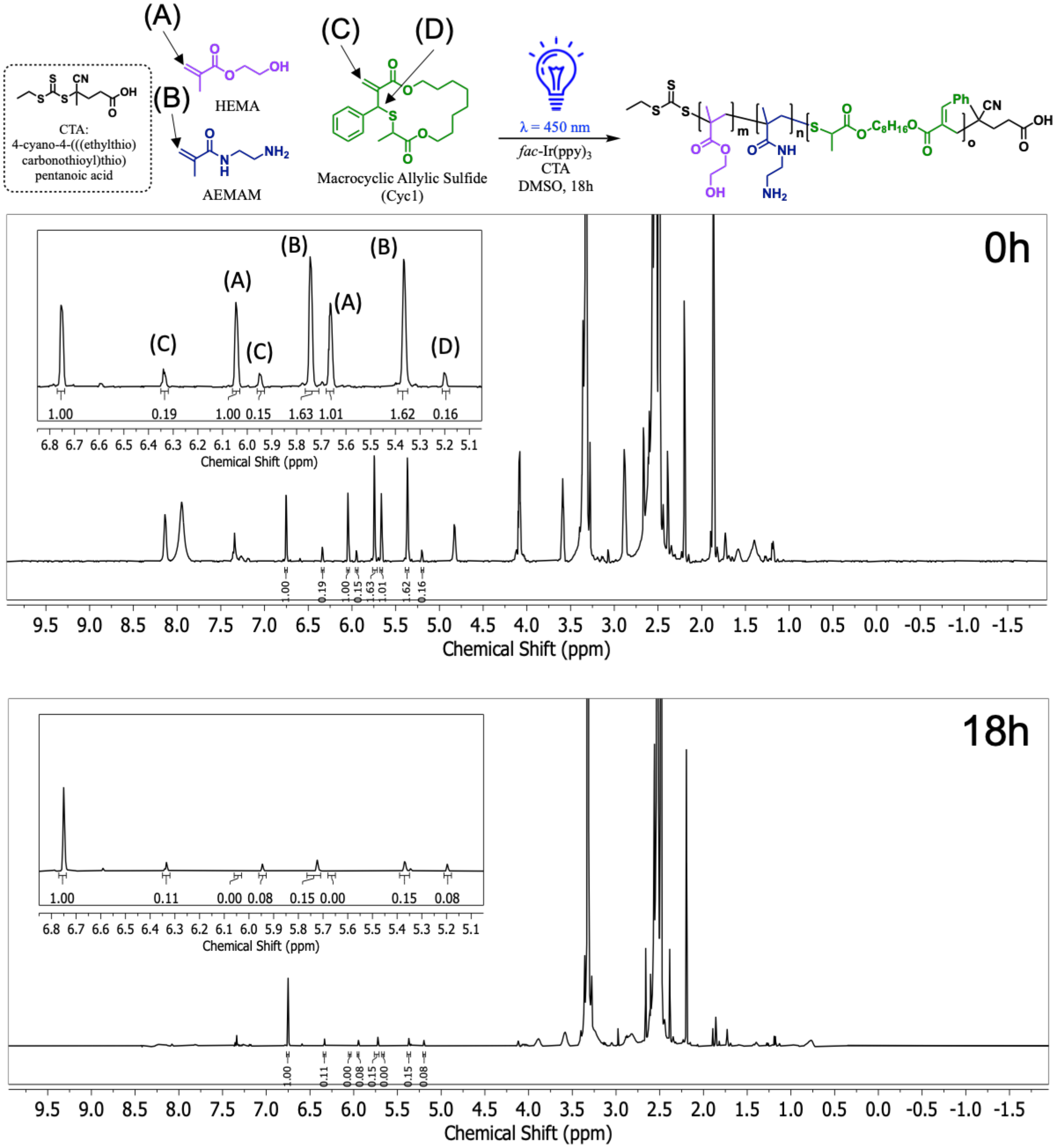


**Figure S8.** [1]H NMR (DMSO-d$_6$) of reaction mixture for degradable cationic copolymer **P2** (*$f_{AEMAm}{}^0$* = 0.6, *$f_{HEMA}{}^0$ = 0.325, $f_{Cyc1}{}^0$ = 0.075)* at the start (0h, top) and end (18h, bottom) of the reaction. Integrals are assigned to vinyl protons of unreacted HEMA (δ = 6.04 ppm and δ = 5.66 ppm, "A"), vinyl protons of unreacted AEMAm (δ = 5.74 ppm and δ = 5.36 ppm, "B"), vinyl protons of unreacted Cyc1 (δ = 6.34 ppm and δ = 5.95 ppm, "C"), and allylic proton of unreacted Cyc1 (δ = 5.20, "D").

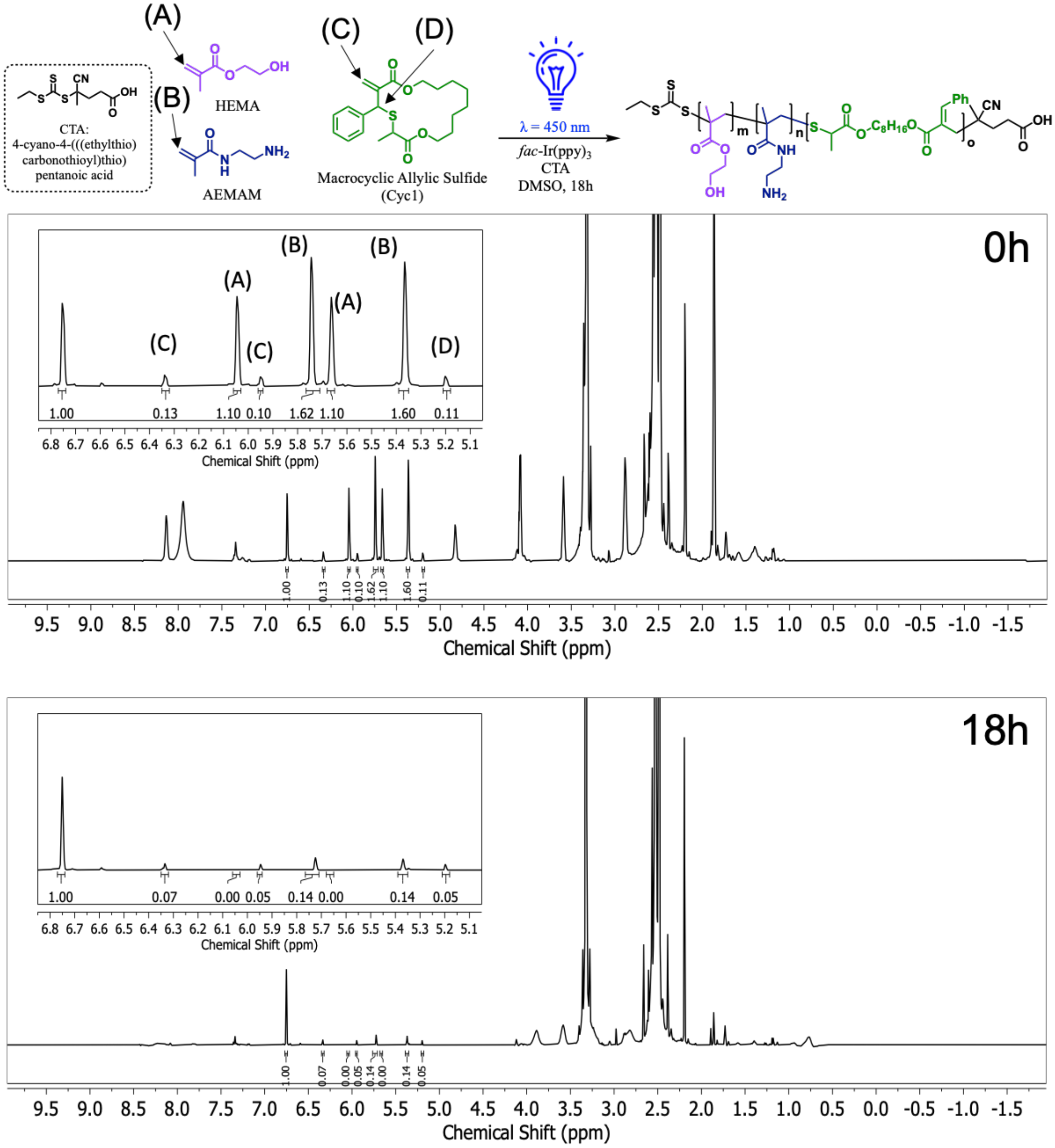


**Figure S9.** [1]H NMR (DMSO-$d_6$) of reaction mixture for degradable cationic copolymer **P3** ($f_{AEMAm}^0$ = 0.6, $f_{HEMA}^0$ = *0.35,* $f_{Cyc1}^0$ = *0.05)* at the start (0h, top) and end (18h, bottom) of the reaction. Integrals are assigned to vinyl protons of unreacted HEMA (δ = 6.04 ppm and δ = 5.66 ppm, “A”), vinyl protons of unreacted AEMAm (δ = 5.74 ppm and δ = 5.36 ppm, “B”), vinyl protons of unreacted Cyc1 (δ = 6.34 ppm and δ = 5.95 ppm, “C”), and allylic proton of unreacted Cyc1 (δ = 5.20, “D”).

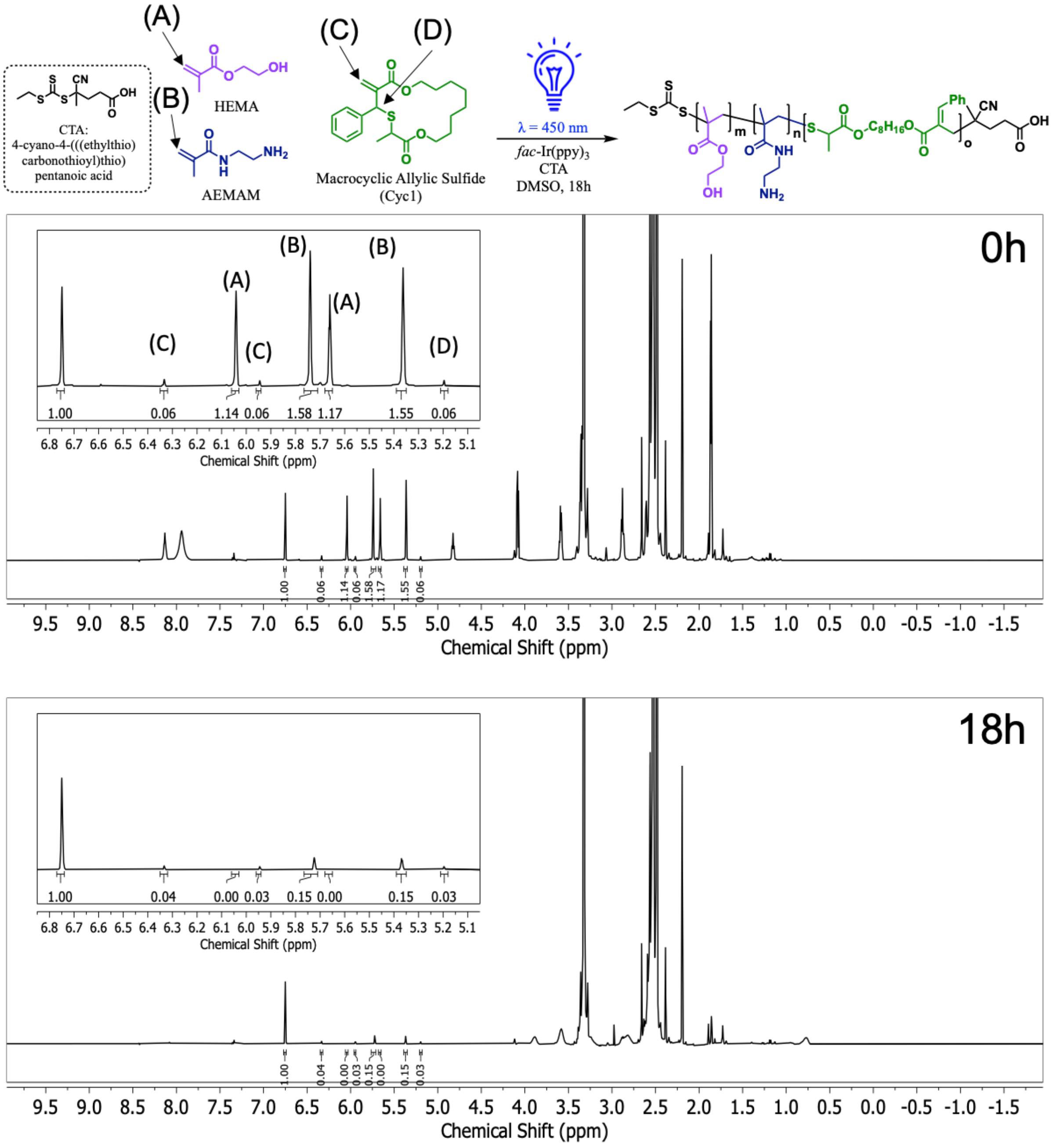

**Figure S10.** $^{1}$H NMR (DMSO-$d_6$) of reaction mixture for degradable cationic copolymer **P4** ($f_{AEMAm}^{0}$ = 0.6, $f_{HEMA}^{0}$ = 0.375, $f_{Cyc1}^{0}$ = 0.025) at the start (0h, top) and end (18h, bottom) of the reaction. Integrals are assigned to vinyl protons of unreacted HEMA (δ = 6.04 ppm and δ = 5.66 ppm, "A"), vinyl protons of unreacted AEMAm (δ = 5.74 ppm and δ = 5.36 ppm, "B"), vinyl protons of unreacted Cyc1 (δ = 6.34 ppm and δ = 5.95 ppm, "C"), and allylic proton of unreacted Cyc1 (δ = 5.20, "D").

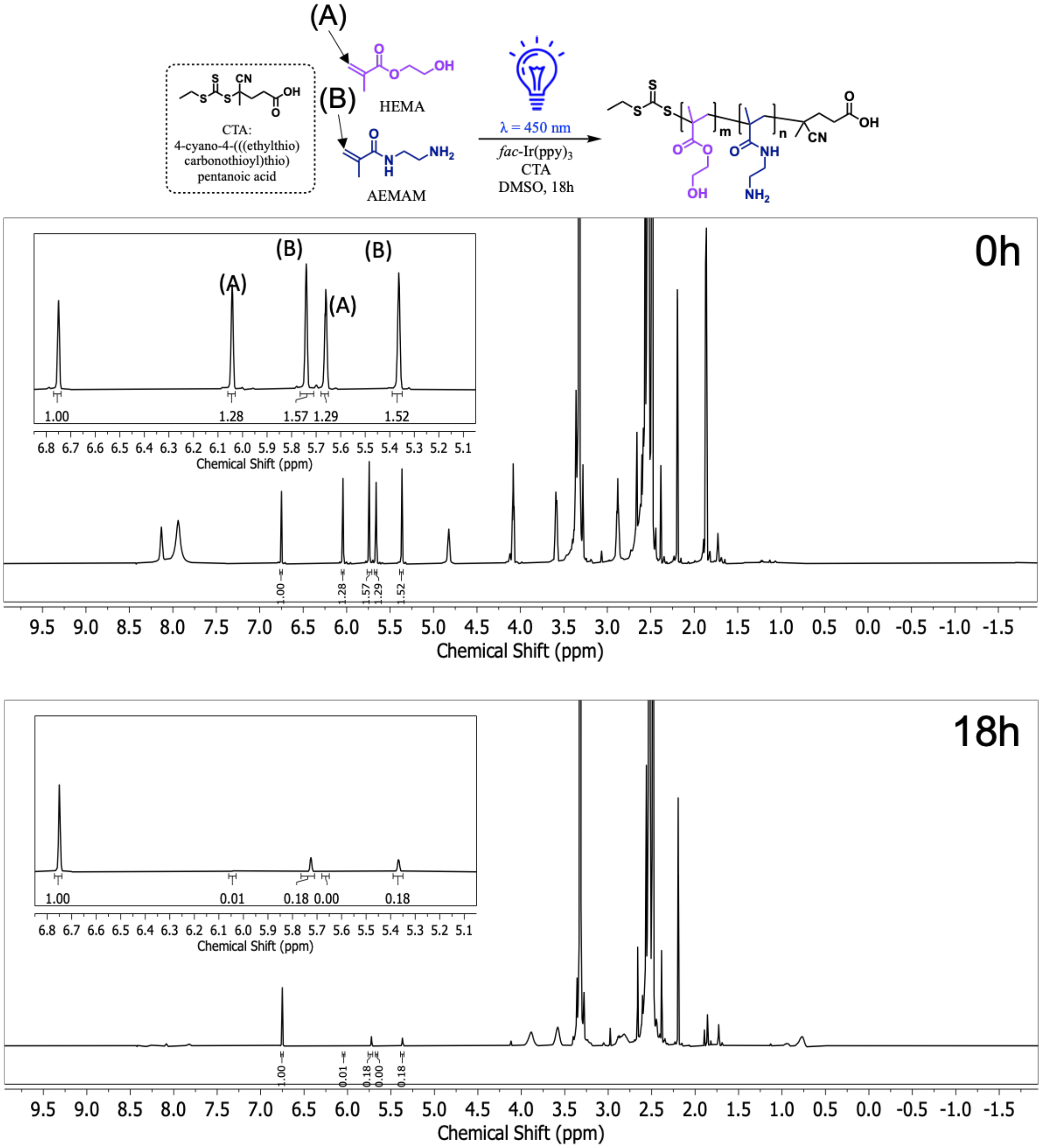


**Figure S11.** $^1$H NMR (DMSO-$d_6$) of reaction mixture for non-degradable cationic copolymer **P5** ($f_{AEMAm}{}^0$ = 0.6, *$f_{HEMA}{}^0$ = 0.4, $f_{Cyc1}{}^0$ = 0)* at the start (0h, top) and end (18h, bottom) of the reaction. Integrals are assigned to vinyl protons of unreacted HEMA (δ = 6.04 ppm and δ = 5.66 ppm, “A”) and vinyl protons of unreacted AEMAm (δ = 5.74 ppm and δ = 5.36 ppm, “B”).

**Table S1.** Summary of monomer conversion of PET-RAFT reactions used to prepare polymer library (**P1-P5**) for *in vitro* transfection and cytotoxicity studies.

| Poly ID | [M]:[CTA] | *Feed Ratio (mol%)* | | | *% Conversion* | | |
|---|---|---|---|---|---|---|---|
| | | HEMA | AEMAm | Cyc1 | HEMA | AEMAM | Cyc1 |
| ***P1*** | 100 | 30 | 60 | 10 | 99.7% | 89.8% | 48.0% |
| ***P2*** | 100 | 32.5 | 60 | 7.5 | 99.8% | 90.9% | 47.0% |
| ***P3*** | 100 | 35 | 60 | 5 | 99.7% | 91.1% | 48.1% |
| ***P4*** | 100 | 37.5 | 60 | 2.5 | 99.7% | 90.5% | 47.3% |
| ***P5*** | 100 | 40 | 60 | - | 99.6% | 88.3% | - |

Monomer conversion was determined based on assigned integrals for vinylic or allylic protons for each of the three unique monomers. Integrals were normalized to mesitylene internal standard ($\delta$ = 6.75 ppm), and change in the normalized integral (100% - $I_{final}/I_{inital}$) was used to calculate monomer conversion. Percent conversion reported for a monomer represents the average change across all integrals assigned to protons for that monomer.

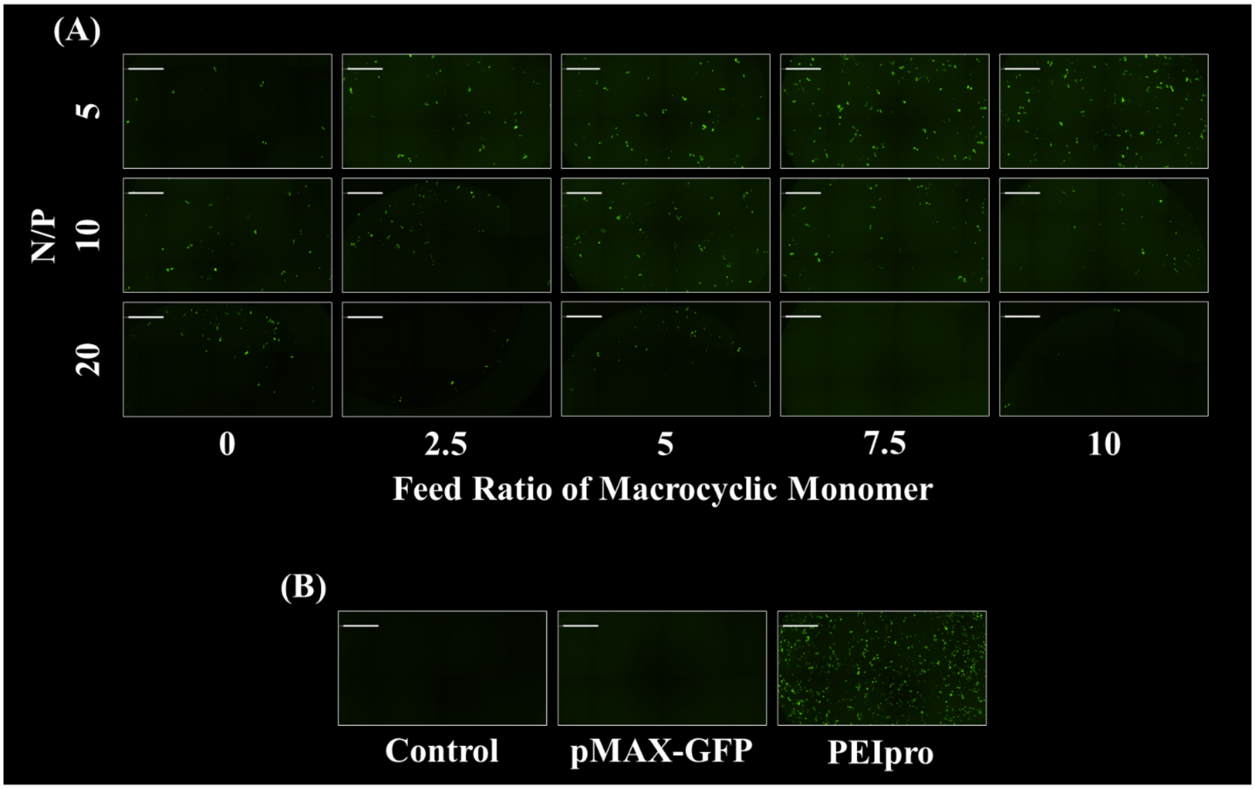


**Figure S12** (A) Fluorescence microscopy of U-2 OS cells transfected with polyplexes formed with degradable and non-degradable polyplexes with varied feed ratio of the degradable residue and varied N/P ratios. (B) Images of U-2 OS control cells, cells transfected only with pMAX_GFP without any polymer delivery vehicle, and cells transfected with PEIpro. GFP expression was measured using target expression analysis with a Celigo Image Cytometer (Nexcelom Bioscience). Green fluorescence channel (483/536) was used to image the GFP expression with an exposure time of 10 ms. Celigo software was used for the automated image analysis that counts the GFP-positive cells and the mean intensity of GFP expression in each cell. Scale bar = 1 mm.
